# Hepatitis C Virus (HCV) diagnosis, epidemiology and access to treatment in a UK cohort

**DOI:** 10.1101/216937

**Authors:** Emily Adland, Gerald Jesuthasan, Louise Downs, Victoria Wharton, Gemma Wilde, Anna McNaughton, Jane Collier, Eleanor Barnes, Paul Klenerman, Monique Andersson, Katie Jeffery, Philippa C. Matthews

## Abstract

**Background:** As direct acting antiviral (DAA) therapy is progressively rolled out for patients with hepatitis C virus (HCV) infection, careful scrutiny of HCV epidemiology, diagnostic testing, and access to care is crucial to underpin improvements in delivery of treatment.

**Methods:** We performed a retrospective study of HCV infection in a UK teaching hospital to evaluate the performance of different diagnostic laboratory tests, to describe the population with active HCV infection, and to determine the proportion of these individuals who access clinical care.

**Results:** Over a total time period of 33 months between 2013 and 2016, we tested 38,510 individuals for HCV infection and confirmed a new diagnosis of active HCV infection (HCV-Ag+ and/or HCV RNA+) in 359 (positive rate 0.9%). Our in-house HCV-Ab test had a positive predictive value of 87% when compared to repeat HCV-Ab testing in a regional reference laboratory, highlighting the potential for false positives to arise based on a single round of antibody-based screening. Of those confirmed Ab-positive, 70% were HCV RNA positive. HCV-Ag screening performed well, with 100% positive predictive value compared to detection of HCV RNA. There was a strong correlation between quantitative HCV-Ag and HCV RNA viral load (p<0.0001). Among the 359 cases of infection, the median age was 37 years, 85% were male, and 36% were in prison. Among 250 infections for which genotype was available, HCV genotype-1 (n=110) and genotype-3 (n=111) accounted for the majority. 117/359 (33%) attended a clinic appointment and 48 (13%) had curative treatment defined as sustained virologic response at 12 weeks (SVR_12_).

**Conclusions:** HCV-Ab tests should be interpreted with caution as an indicator of population prevalence of HCV infection, both as a result of the detection of individuals who have cleared infection and due to false positive test results. We demonstrate that active HCV infection is over-represented among men and in the prison population. A minority of patients with a diagnosis of HCV infection access clinical care and therapy; enhanced efforts are required to target diagnosis and providing linkage to clinical care within high risk populations.

**ABBREVIATIONS:**

DAA: Direct Acting Antiviral
ELISA: Enzyme linked immunosorbent assay
HCV: Hepatitis C Virus
HCV-Ab: IgG antibody to Hepatitis C virus
HCV-Ag: Hepatitis C virus core antigen
HCV RNA: Hepatitis C ribonucleic acid (viral load)
MSM: men who have sex with men
NAT: nucleic acid testing
PCR: polymerase chain reaction (test for viral load)
PPV: positive predictive value
PWID: people who inject drugs
SDG: Sustainable Development Goals
SVR: sustained virologic response
WHO: World Health Organisation

## INTRODUCTION

The World Health Organization (WHO) estimates that 71 million people are chronically infected with the Hepatitis C Virus (HCV), and 0.4 million people die each year as a consequence [1, 2]. United Nations Sustainable Development Goals (SDGs) [3] and Global Health Sector Strategy on Viral Hepatitis [2] have a target of elimination of viral hepatitis as a public health threat by the year 2030. The need for enhancing HCV diagnosis has acutely become more pertinent as a result of the availability of highly effective Direct Acting Antiviral (DAA) treatment [4]. On a worldwide basis, an estimated 80–85% of individuals with chronic infection are unaware of their infection [5, 6], although this figure may be somewhat lower in the UK, and among those with a diagnosis a minority currently receive treatment.

Enhanced diagnosis of HCV is important not only as a pathway to treatment for individual patients, but also to allow confident estimates of the true prevalence of chronic HCV in different settings, to underpin appropriate allocation of resources and development of infra-structure for treatment [7]. Diagnosis of HCV is based on three different approaches to testing, which may be used alone or in combination. These are (i) detection of an IgG antibody by ELISA (HCV-Ab); (ii) detection of HCV core antigen (HCV-Ag); (iii) Nucleic acid testing (NAT) to detect HCV RNA by PCR (Table 1)

**Table 1:** Comparison of diagnostic laboratory tests used to detect exposure and activity of HCV infection

| Screening tool | HCV-Ab | HCV-Ag | PCR for HCV RNA |
| --- | --- | --- | --- |
| Pro's | ♦ Widely available<br>♦ Inexpensive<br>♦ Much experience and data for use as first-line approach to screening for HCV exposure (underpins many old seroprevalence studies) | ♦ Diagnostic of active infection (not past exposure)<br>♦ Improved specificity and reduced window period compared to HCV-Ab [8-13]. | ♦ Accepted gold-standard diagnostic test for active infection (not past exposure)<br>♦ Allows quantitative monitoring of viraemia; useful for monitoring therapy<br>♦ Genome amplification allows other information to be ascertained (e.g. genotype; drug resistance).<br>♦ Can potentially be applied to dried blood spots (DBS) |
| Con's | ♦ Subject to inter-assay variability and a variable rate of false positive results [14, 15]; false positive has been associated with ethnicity [16, 17], age [16], raised IgM and erythrocyte sedimentation rate (ESR) [14], auto-antibodies [18], and prosthetic devices [19].<br>♦ Test of exposure, not of active infection | ♦ Not universally available<br>♦ More expensive than HCV-Ab<br>♦ Not consistently regarded as sufficiently sensitive to replace PCR | ♦ Not universally available<br>♦ Expensive: beyond the financial reach of many resource-limited settings |

Reliance upon HCV-Ab screening has potentially distorted epidemiological data upon which resource-planning depends [20], as detection of individuals who have cleared infection, as well as false positives, may have led to over estimation of HCV prevalence. As a result, there has been a progressive move towards using HCV-Ag and/or HCV PCR to determine accurately the population prevalence of active infection [1]. Although sensitivity and specificity of HCV-Ag testing appears to perform well when compared head-to-head with PCR [7, 13], there are still potential doubts over whether this test is adequately sensitive to be widely implemented as a primary screening tool, and recent WHO guidelines continue to recommend use of the HCV-Ab test as first line [1]. A careful balance must be struck between managing cost and optimising specificity without sacrificing sensitivity [21–24].

We here set out to assess our progress in diagnosing and treating HCV infection in a tertiary referral UK hospital, in the context of global aspirations for elimination. We reviewed the performance of our local HCV testing protocol in two different time periods, first when screening was undertaken using an HCV-Ab test only, and subsequently following the introduction of a combined approach using both HCV-Ab and HCV-Ag testing. In each case, we went on to evaluate further using HCV PCR. Collating these data allowed us to evaluate the performance of different diagnostic tests, to describe the epidemiology of our local cohort, and to determine the proportion of those with active HCV infection who attend a hepatology clinic and receive treatment.

## METHODS

### Setting and cohort

Our microbiology laboratory serves a large UK tertiary referral and teaching centre (http://www.ouh.nhs.uk/) that provides one million patient contacts a year, and handles samples referred from the community as well as four in-patient sites. We retrospectively interrogated electronic microbiology records for all HCV screening tests performed within two defined time-intervals, during which different diagnostic algorithms were operating. These are summarized in Fig 1 and outlined as follows:

**Figure 1:**
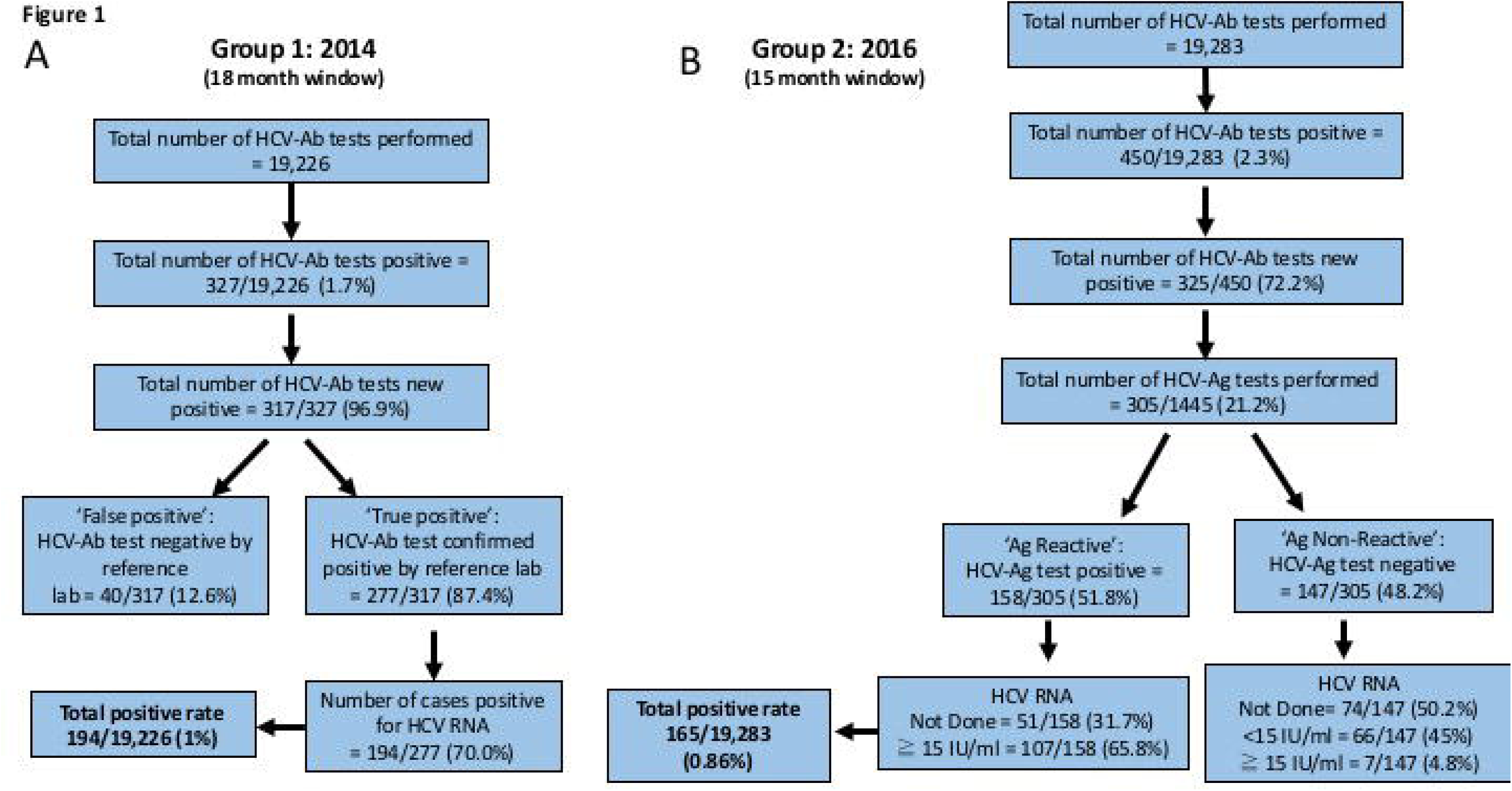
Algorithms describing approach to, and results of, HCV diagnostic testing in a UK teaching hospital laboratory in 2014 (A) and 2016 (B).

i. **Group 1** (18 months; January 2013 - June 2014; Fig 1A). During this period, samples were screened for HCV-Ab using an ADVIA Centaur automated immunoassay (Bayer). HCV-Ab-positive samples (excluding repeat samples from patients with a pre-existing HCV diagnosis) were sent for confirmatory testing by the regional reference laboratory (Public Health England, Colindale), using two further ELISA tests (Ortho and BioRad). Antibody positive samples (based on sample:cut-off ratio >1) were tested for HCV RNA using Abbott HCV M2000 assay.
ii. **Group 2** (15 months; January 2015 - March 2016; Fig 1B). HCV diagnosis during this period was undertaken using a combination of HCV-Ab and HCV-Ag testing using Abbott Architect i2000SR with Diasorin Liason XL for confirmation of HCV-Ab. HCV-Ag assay positives are in the range 10-20,000 fmol/L.

For those testing positive with the initial screening test, we then collected follow-up testing data (per algorithms in Fig 1). We recorded patient age, sex, and the location from which the sample was sent. Treatment data were captured and recorded from an electronic database within the Hepatology Department. Response to treatment was defined as SVR_12_ (sustained virologic response at ≥12 weeks following the end of therapy).

Ethnic origin is not routinely captured data in hospital electronic systems. Prior to anonymisation, we therefore used an analytical tool to estimate ethnicity, applying Onolytics software for all patients for whom a full name was part of the electronic record (http://onolytics.com [25–27]). This software was developed in 2006-7 funded by Economic and Social Research Council (ESRC) Knowledge Transfer Partnerships, and sets out to determine probable ethnic origin based on name.

### Data analysis

Statistical analysis was performed using GraphPad Prism v.7.0b and Googlesheets (https://docs.google.com/spreadsheets). We compared binary values using Fisher’s Exact Test, Mann-Whitney U test for continuous non-parametric data and linear regression for correlation between continuous variables.

### Ethics approval

Ethics approval was not required, as this study was undertaken as a departmental quality improvement exercise within microbiology using anonymised patient data, and completed the audit cycle for previously approved audit projects [28, 29]. Data for Onolytics analysis were handled separately and were subject to a confidential disclosure agreement drawn up by University of Oxford Research Services (February 2016).

## RESULTS

### HCV testing: prevalence and characteristics of infection

In total, we present data for 38,510 HCV tests done during the two intervals reviewed. On average testing increased slightly, from an average of 1068 tests / month during the earlier phase of the study (Group 1) up to 1286 tests / month in the later time period (Group 2); (Fig 1).

In total, 359 new active HCV infections were identified and confirmed during the two testing periods (359 / 38,510 = 0.9% of all tests performed; Fig 1). Characteristics of these individuals are summarised in Table 2 and the complete metadata are available on-line (https://doi.org/10.6084/m9.figshare.5355097.v2). The median age was 37 years (IQR 31-48) and there was a consistent male excess, with men representing 55% of those tested [28] but 85% of all new diagnoses (Table 2). Over one-third (36%) of new diagnoses were made in prison. Of the remainder, the majority were in primary care or attending hospital outpatient departments.

**Table 2:** Characteristics of individuals with active HCV infection in a UK teaching hospital in two time windows between 2014 and 2016

|  | Group 1<br>(2014) | Group 2<br>(2016) | Group 1 +<br>Group 2<br>(2014-2016) |
| --- | --- | --- | --- |
| Total number confirmed positive for active HCV infection | 194 | 165 | 359 |
| Number male (% of positive tests) | 167 (86.1%) | 137 (83.0%) | 304 (84.7%) |
| Age in years (median and IQR) | 39 (31-49) | 37 (30-44) | 37 (31-48) |
| Location<br>- Prison<br>- Hospital in-patient<br>- Hospital out-patient / primary care<br>- Sexual health clinic<br>- Occupational health<br>- Primary care for homeless (drug and alcohol service) | 63 (32.5%)<br>15 (7.7%)<br>90 (46.4%)<br>16 (8.2%)<br>2 (1%)<br>8 (4.2%) | 66 (40.7%)<br>13 (8%)<br>64 (38.8%)<br>13 (7.9%)<br>1 (0.6%)<br>8 (4.8%) | 129 (36.2%)<br>28 (7.9%)<br>154 (42.9%)<br>29 (8.1%)<br>3 (0.8%)<br>16 (4.5%) |
| Ethnic origin<br>- Black<br>- Asian<br>- European<br>- Unknown | 7 (3.6%)<br>18 (9.3%)<br>149 (76.8%)<br>20 (10.3%) | 1 (0.6%)<br>7 (4.2%)<br>97 (60.7%)<br>57 (34.5%) | 8 (2.2%)<br>25 (7.0%)<br>246 (69.4%)<br>77 (21.4%) |

Numbers of positive tests are shown with percentage of positive tests in brackets. Median age for each group is shown with IQR in brackets.

Genotype was available in 250 cases (70% of new diagnoses), with genotypes 1 and 3 accounting for the majority (44% each; Fig. 2). This is in keeping with the overall genotype distribution reported for Europe [30], and Public Health England data (90% of all cases accounted for by genotypes 1 and 3) [31].

**Figure 2:**
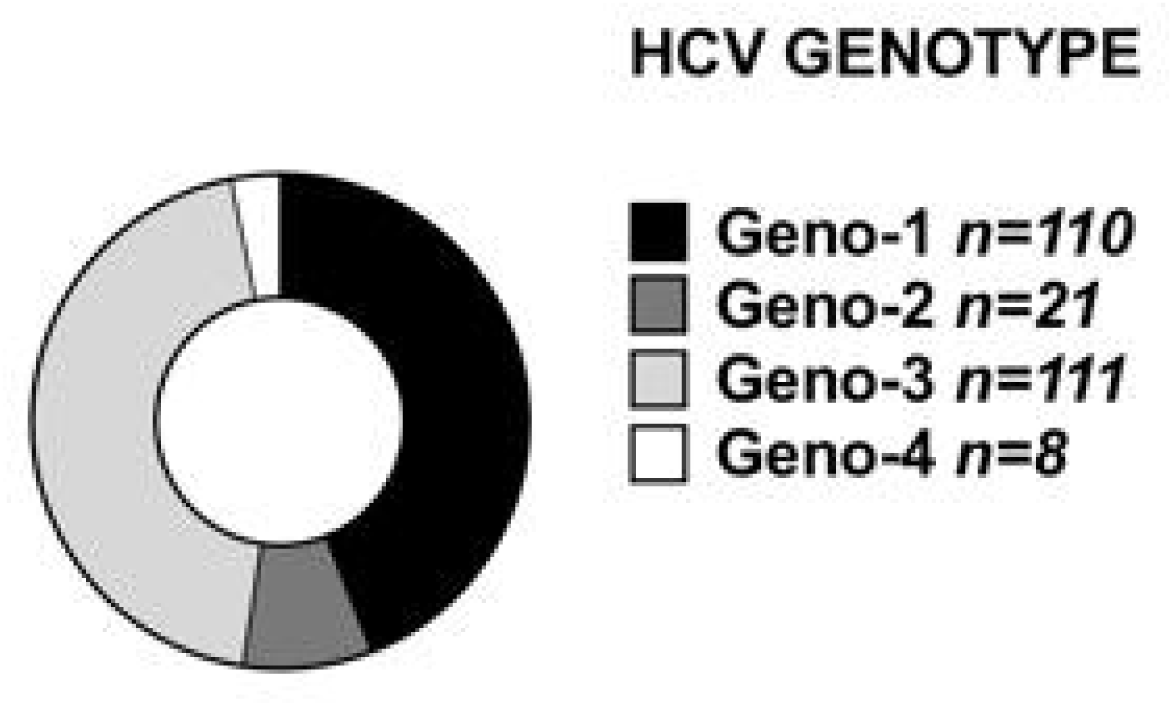
Distribution of HCV genotypes in a UK cohort. Data for 250 individuals with a new laboratory diagnosis of HCV infection are shown. There was no enrichment of a specific genotype in the prison population (prison population accounted for 30/84 geno-1 infections, and 34/80 geno-3 infections; p=0.4 Fisher’s Exact Test).

### HCV-Ab test outcomes and performance

In the earlier testing period (Group 1), 277 of 317 HCV-Ab positive samples were positive on confirmatory testing for HCV-Ab at the reference laboratory (Fig 1A), giving our in-house test a positive predictive value (PPV) of 87.4% compared to a regionally accepted standard. We used these results to investigate whether any host factors are associated with false positive antibody tests, and found that individuals identified as African have a higher chance of a false-positive HCV Ab test (Fig 3). We confirmed this result by multivariate logistic regression analysis, in which African ethnicity was significantly associated with a false positive Ab test result (p=0.0004), but age >60 years and gender were not. Prison location was associated with a true positive Ab-test result (p=0.01).

**Fig 3:**
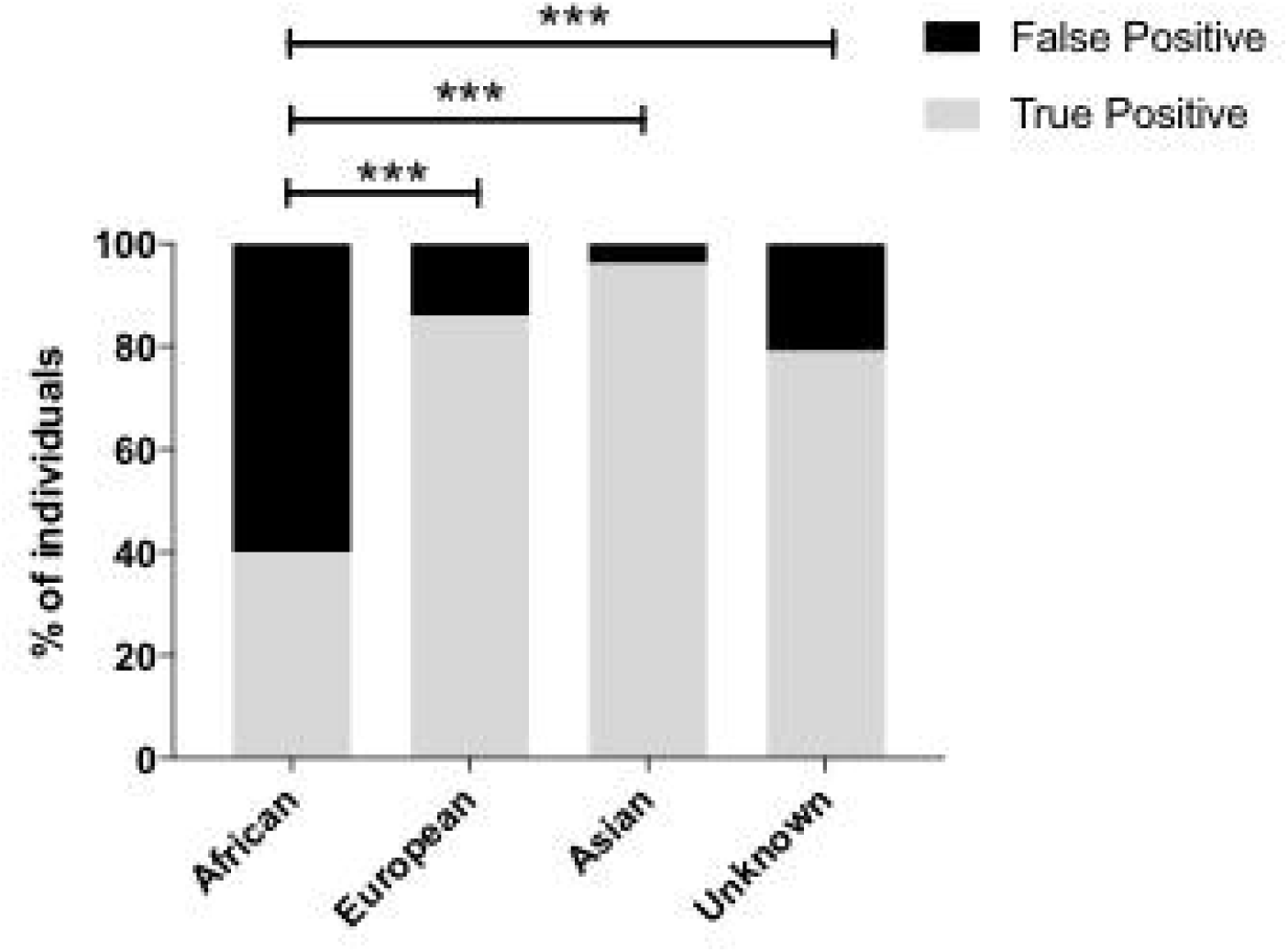
False positive HCV IgG antibody results according to ethnic origin in a UK cohort. Ethnicity was estimated using Onolytics software [26, 27]. Data shown are for a cohort recruited starting in 2014, screened using an inhouse HCV-Ab (ADVIA Centaur automated immunoassay; Bayer) and confirmed using two further ELISA tests (Ortho and BioRad). ‘False positives’ are defined as those screening positive on ADVIA and negative on confirmation, ‘true positives’ are defined as samples positive on all three tests. P-values obtained by Fishers Exact Test shown as * p<0.05, ** p<0.005, *** p<0.0005.

### HCV-Ag test outcomes and performance

In the later testing period (Group 2), the PPV of the combined use of HCV-Ab plus HCV-Ag was 100% when compared to a gold-standard diagnostic test using PCR (Table 3). This exceeds a previous estimation of the PPV of the Ag test (94.7%) calculated from assimilation of data from other comparable reports [13].

**Table 3:**
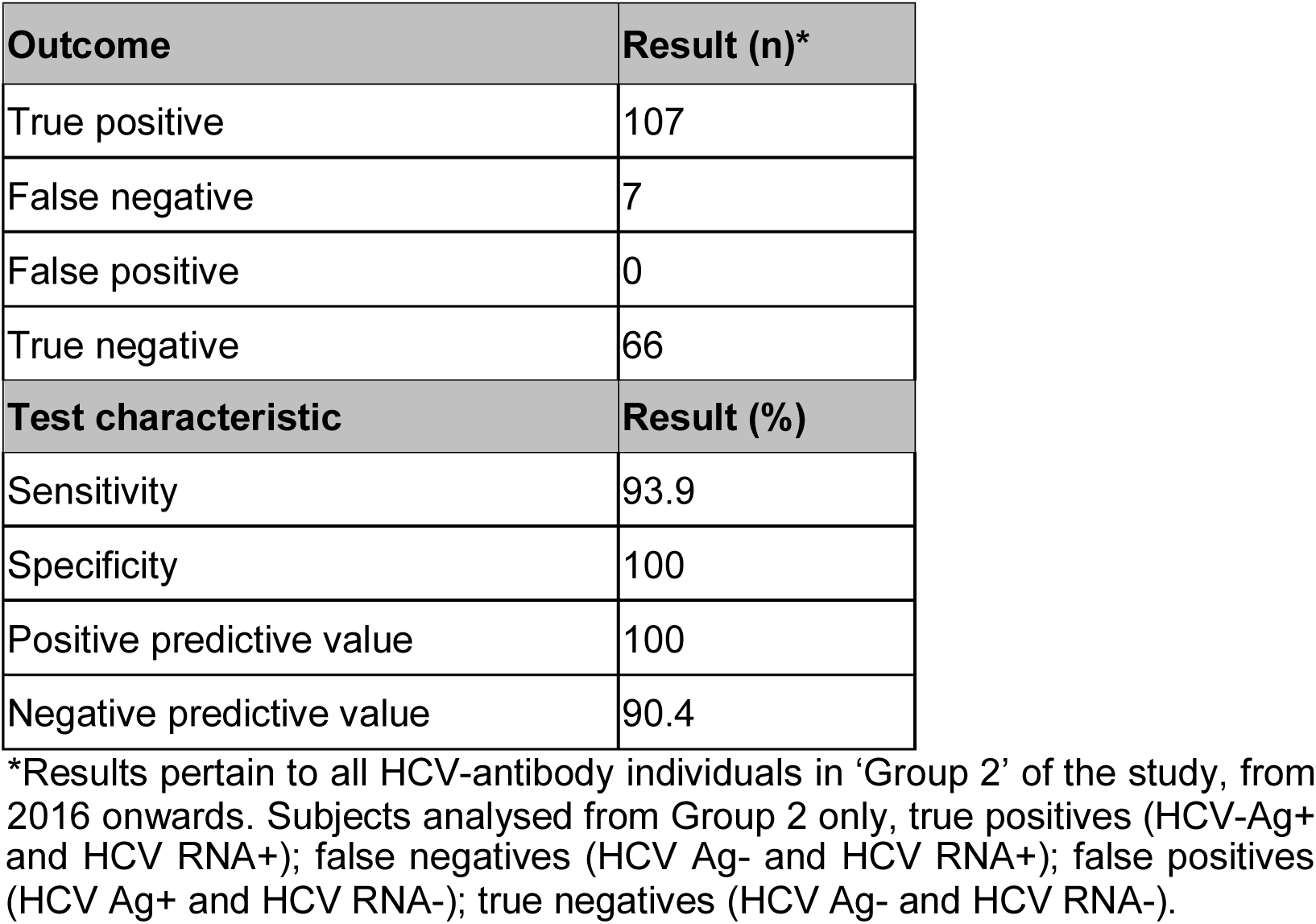
Outcome of diagnostic testing for HCV infection using core antigen detection (HCV-Ag) compared to gold standard PCR for HCV RNA.

| Outcome | Result (n)* |
| --- | --- |
| True positive | 107 |
| False negative | 7 |
| False positive | 0 |
| True negative | 66 |
| Test characteristic | Result (%) |
| Sensitivity | 93.9 |
| Specificity | 100 |
| Positive predictive value | 100 |
| Negative predictive value | 90.4 |

\*Results pertain to all HCV-antibody individuals in 'Group 2' of the study, from 2016 onwards. Subjects analysed from Group 2 only, true positives (HCV-Ag+
and HCV RNA+); false negatives (HCV Ag- and HCV RNA+); false positives (HCV Ag+ and HCV RNA-); true negatives (HCV Ag- and HCV RNA-).

We explored the relationship between HCV-Ag and HCV RNA. Individuals with a positive HCV-Ag test had a median HCV viral load of 6.3 × 10^5^ IU/ml (Fig 4A), and there was a significant positive correlation between quantitative antigenaemia and viral load (r=0.67, p<0.0001; Fig 4B). This suggests that, in the absence of having access to a quantitative PCR result, HCV-Ag is a good surrogate marker of viraemia. However, in a small proportion of cases, the antigen test results in false negative results (Table 4). There are no consistent features that unify these misleading HCV-Ag results; in particular there was no clear relationship between false negative HCV-Ag and low viral load (although one sample had HCV RNA 25 IU/ml).

**Figure 4:**
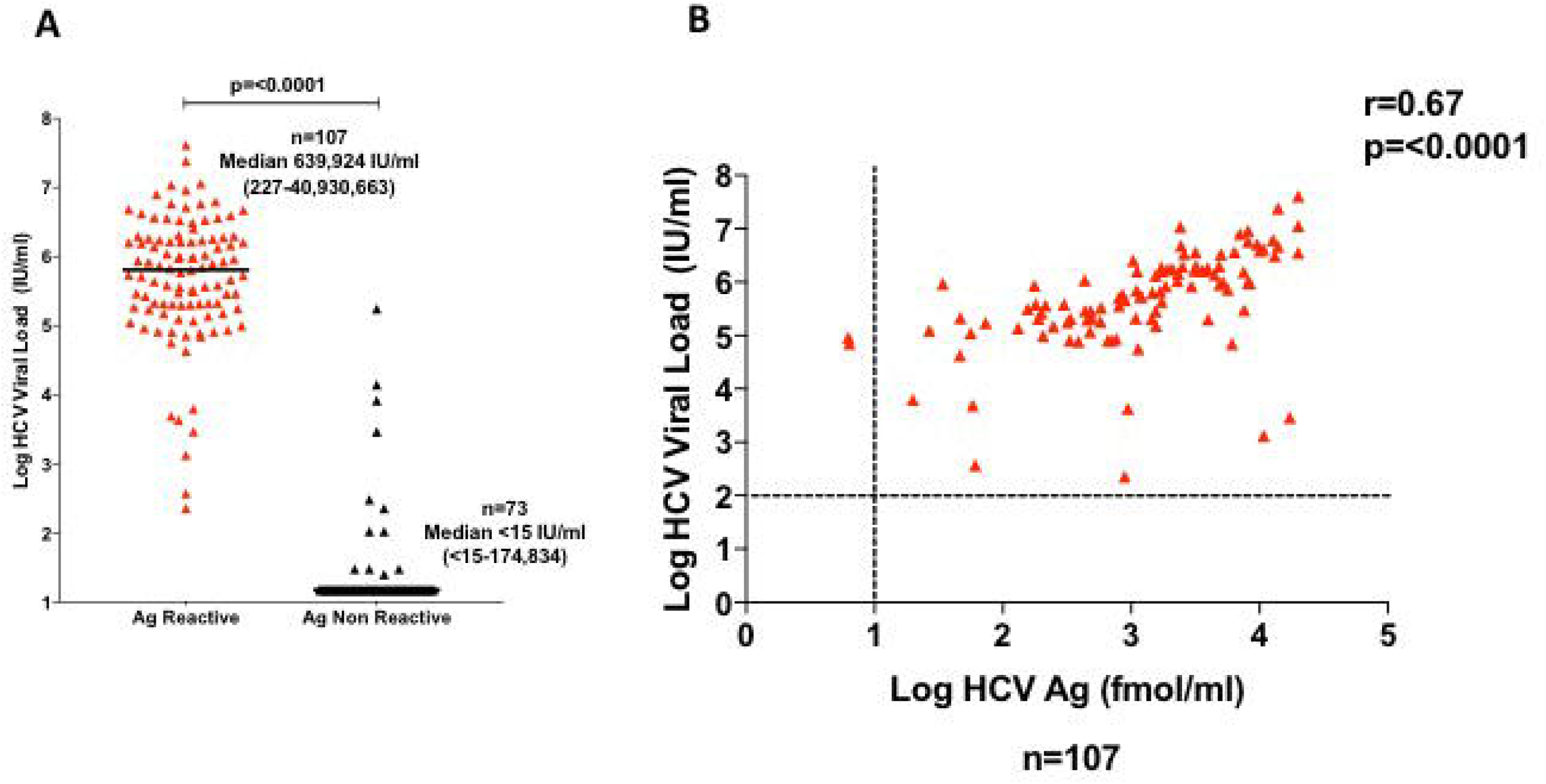
Relationship between HCV Antigen test and quantitative PCR for HCV RNA viral load. (A) Range of HCV viral loads for samples testing HCV-Ag positive (n=107) and HCV-Ag negative (n=73). (B) Linear regression plot showing correlation between HCV-Ag and HCV-RNA for all samples testing HCV-Ag positive (n=108). Dashed lines represent threshold for detection for HCV RNA (15 IU/ml) and HCV-Ag (10 fmol/ml).

**Table 4.** Summary of patients who tested false negative for HCV core antigen, using HCV RNA PCR as a gold-standard reference test.

| Age group (years) | Sex | Patient location | Ethnicity | HIV status | HCV Ag (fmol/ml) | HCV Ab (sample/cut-off ratio) | Geno-type | HCV viral load (IU/ml ) |
| --- | --- | --- | --- | --- | --- | --- | --- | --- |
| 30-39 | F | Sexual health | Unknown | negative | 0.0 | 11.8 | IS | 25 |
| 40-49 | M | GP | European | negative | 0.0 | 12.2 | NR | 226 |
| 50-59 | F | Medicine | European | yes | 0.0 | 3.1 | IS | 302 |
| 20-29 | M | Prison | European | negative | 0.62 | 14.2 | 2b | 2916 |
| 20-29 | M | Prison | European | NR | 0.00 | 12.6 | NR | 8232 |
| 30-39 | F | Out-patient | European | NR | 1.98 | 15.8 | 1b | 13860 |
| 30-39 | M | Prison | European | negative | 0.00 | 12.2 | 3a | 174834 |

NR = not requested; IS = insufficient sample. Total number of HCV core antigen tests carried out in this period n=306. False negative subject (n=7) HCV RNA viral loads did not differ significantly to others ( $p=0.187$ Mann Whitney U test). None of the
patients with a false negative result underwent a repeat Ag test so laboratory error cannot be ruled out in this instance.

Of 359 patients with a new HCV diagnosis, 117 (33%) attended a hepatology clinic appointment, 76 were treated (21%) and 48 had an outcome of SVR_12_ (13%). These data illustrate the substantial loss of patients at each step of the clinical pathway (Fig 5; Table 5), due to a combination of factors including poor linkage between services, itinerant populations, individuals who are drug users and/or in prison, and deaths.

**Figure 5:**
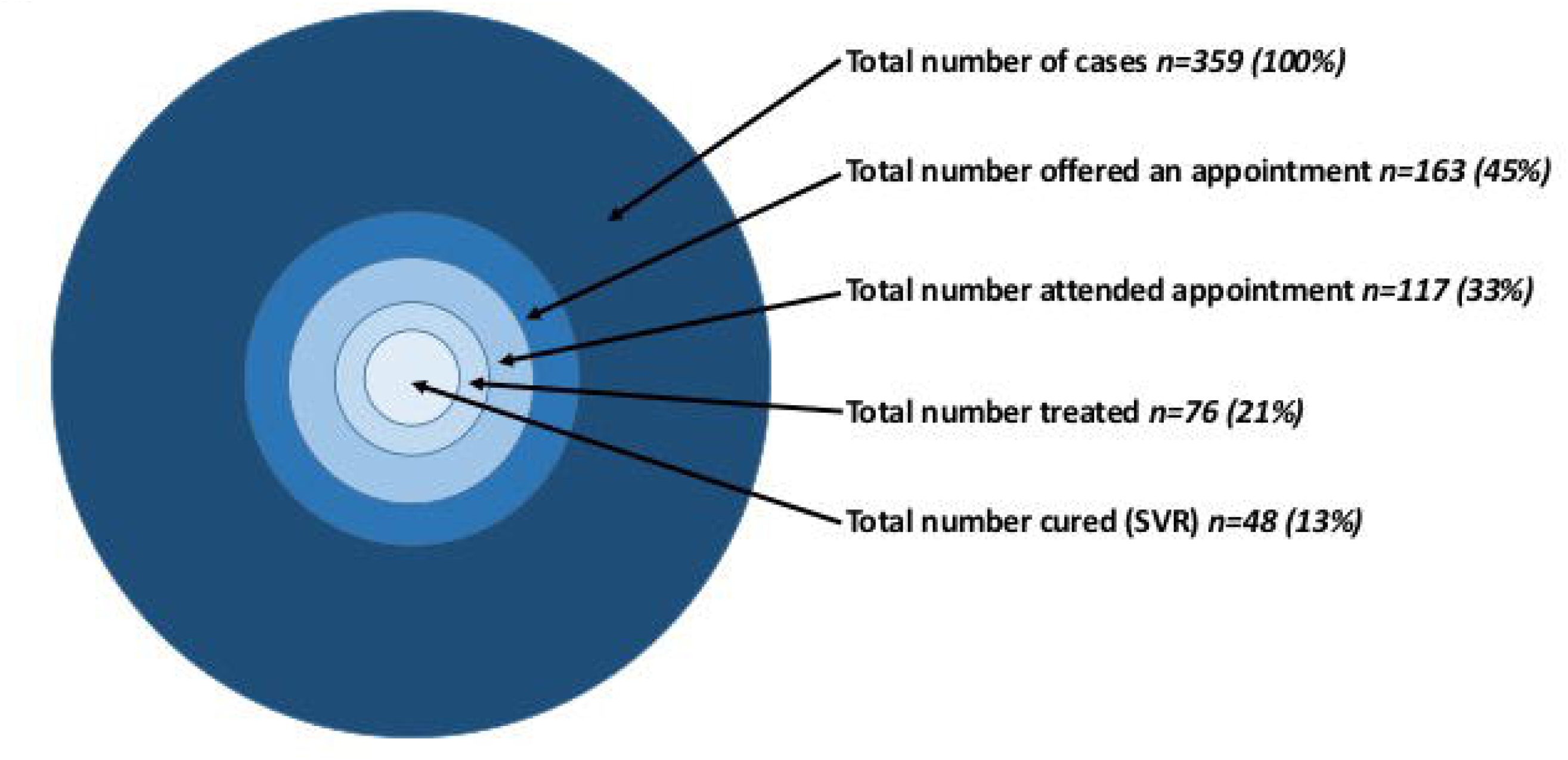
Graphical representation of the disparity between the number of individuals diagnosed with active HCV infection and those who access clinical review, treatment, and achieve SVR Summary of outcomes for the entire cohort is shown in table 4. The percentages quoted in this figure represent the proportion of patients in each category from the total denominator of 359. Among those with a known treatment outcome, 30/129 (23%) of those from prison attended a clinic appointment, while in the nonprison populations 160/230 (70%) have already been seen or have an appointment to be seen in clinic (p<0.0001, Fisher’s Exact Test).

**Table 5:** Summary of clinical care outcomes in 359 patients with a diagnosis of chronic HCV infection

| Treatment Status | Patient Classification | Number (%) |
| --- | --- | --- |
| **Not treated** | Offered appointment but did not attend | 46 (12.8) |
|  | Seen in sexual health clinic | 28 (7.8) |
|  | Seen by hepatology but not yet treated | 24 (6.7) |
|  | Died | 11 (3.1) |
|  | Transferred out of area | 10 (2.8) |
|  | Spontaneous clearer | 3 (0.8) |
| **Treated or awaiting treatment** | Treated - sustained virologic response at ≥12 weeks (SVR<sub>12</sub>) | 47 (13.1) |
|  | Treated - no outcome data | 20 (5.6) |
|  | Awaiting treatment | 9 (2.5) |
|  | Treated – relapsed | 8 (2.2) |
| **Unknown** | No data / not known in Oxford | 149` (41.5) |
|  | Prison clinic | 3 (0.8) |
|  | Paediatric clinic | 1 (0.3) |
| **TOTAL** |  | 359 (100) |

This study was not designed to examine or report on the outcomes of treatment. However, we examined existing treatment data (Suppl data table) to look for evidence of different outcomes between genotypes 1 and 3. Among genotype 1 infections, we recorded 28 cases of SVR_12_, and two cases who relapse. For genotype 3, there were 16 cases of SVR_12_ and five relapses. This difference did not reach statistical significance (p=0.1, Fisher’s Exact Test).

## DISCUSSION

### Summary comments

Careful scrutiny of HCV testing and treatment is important so that best efforts can be made to diagnose and treat individuals with active infection in order to optimize the benefits of recent therapeutic advances, and to move towards United Nations Sustainable Development Goals [3, 6]. Our study indicates a high workload of HCV-screening in a UK teaching hospital laboratory (>1000 tests performed each month). Overall, 0.9% of these were confirmed to have active HCV infection, with a substantial excess among males, and over one-third from the prison population. Following the implementation of HCV-Ag testing as part of the diagnostic algorithm, the PPV of a positive test increased to 100%, slightly exceeding that reported by other recently published studies [13]. In keeping with national and international data, genotypes 1 and 3 predominate [30, 31], with a trend towards better outcome in treatment of genotype 1 infection.

### Relevance to laboratory and clinical practice

Although HCV-Ag testing can potentially replace a nucleic acid test for HCV diagnosis or monitoring in some settings [13, 32, 33], guidelines from the UK [9], North America [34] and the WHO [1] still advocate use of PCR as a definitive test following HCV-Ab (± HCV-Ag) screening. RNA PCR also remains the gold-standard approach to monitoring progress during and after treatment and is required for genotyping which is currently still important to underpin optimum choice of DAA regimen, despite an ultimate desire to develop pan-genotype treatment [35]. Ultimately, therefore, reducing the cost of PCR may be a more desirable outcome than focusing on improving the sensitivity of antigen detection.

The small proportion of all diagnosed patients who access clinical care and receive successful treatment is in keeping with that reported in other centres including recently by a London centre [36], reflecting many challenges for HCV elimination. Our data highlight the particular attention that is needed for the vulnerable prison population; with a worldwide estimate of 15% HCV prevalence in prisoners worldwide [37], this is an issue that demands urgent international attention. Offering treatment within the prison system has now become a realistic possibility, on the basis of oral DAA therapy, shortened treatment regimens (regimens of 12 to 24-weeks, and possibly shorter), and a low rate of side-effects [38].

In the longer term, bigger datasets are required to improve our insights into this patient population, and to identify areas where additional resource and investment is required. Substantial efforts are underway, with funding underpinned by the UK National Institute for Health Research (NIHR) through the NHIR Health Informatics collaborative (NHIC), to improve the collation of clinical and laboratory data for patients with viral hepatitis, in order to develop and strengthen links between patient care, laboratory microbiology, and research questions [36].

### Significance of male excess

There are emerging data to suggest a genuine discrepancy in susceptibility to, and outcomes of, infection between males and females [39, 40], but in this instance the male predominance may be accounted for by behaviour (e.g. among MSM and PWID) rather than biological differences. In the context of this study we do not have the careful prospective socio-demographic data that would be required to investigate this further.

### Caveats and limitations

Our analysis must be set in the overall context of the low prevalence of HCV in our setting, and the retrospective approach to data collection. Such approaches to estimation of epidemiology underestimate the true prevalence, as a result of a large pool of individuals who are HCV-infected but never receive a test [6].

The rate of false negative HCV tests is our population is likely to be low, but quantifying this was not possible within this study, as we relied on identifying samples that initially tested positive. In order to ascertain the PPV of the HCV-Ab test in-house, we referred to a Reference Laboratory test as ‘gold standard’. However, this repeat testing in a Reference Laboratory setting is itself subject to an error rate, and therefore may lead to a misrepresentation of our overall assay performance.

We found evidence that the HCV-Ab test performs poorly in individuals predicted to be of African origin; similarly, a similar high rate of false positive tests has previously been reported from Polynesia [41]. This illustrates how tests that have been developed and tested in white European/Caucasian populations cannot necessarily be robustly applied in other settings, potentially contributing to global healthcare inequalities. Although name can be a reliable surrogate for ethnic background, and the tools used here have been validated [26, 27], this remains an imperfect way to derive ethnic origin and is potentially confounded by a variety of factors, the most obvious of which is individuals who change their name (usually in the setting of marriage).

In our setting, the sexual health clinic anonymises patient data which prevents robust linkage between services, and we are unable to trace outcomes for patients who were diagnosed via this route (8% of the total; Table 2). Likewise, consistent identification and tracing of individuals who are drug users and/or in prison is challenging, and we cannot exclude the possibility of duplication of some of these individuals within our dataset.

### Performance of HCV-Ag test

We identified seven patients in this cohort in whom there was discordance between HCV-Ag (negative) and HCV RNA results (positive). There was no consistent feature (age, sex, patient location, genotype of HCV infection) that unified these individuals (Table 4). One potential explanation for discrepancies is mutations in the core region of the HCV genome which could account for a failure of antigen detection [42], or potentially cause lack of PCR amplification if mutations occur in the regions required for primer binding.

### Global context

Moving towards the SDG target [2] requires a multi-faceted approach, with focus on optimization of laboratory testing and reduction of costs in order to improve access to accurate diagnosis, advocacy for better testing and treatment for populations in resource-limited settings, allocating resources in order to deliver curative therapy supported by appropriate monitoring, targeting interventions at high-risk populations including MSM, PWID and prisons, and ensuring that individuals diagnosed with infection are offered – and receive – clinical care and follow-up.

## DECLARATIONS

### Ethics approval and consent to participate

No specific ethics approval was required for this study as it was undertaken as a combination of audit and quality improvement from within a clinical microbiology laboratory and hepatology service; data were anonymised prior to analysis and no interventions were implemented.

### Consent for publication

Not applicable.

### Availability of data and material

The datasets generated and/or analysed during the current study are available in the Figshare repository [https://doi.org/10.6084/m9.figshare.5355097.v1].

## Authors’ contributions

Study conception and design: PK, KJ, PCM. Data collection: GJ, LD, VW, GW. Running and interpreting clinical laboratory tests: GJ, MA, KJ, PCM. Data analysis: EA, AM, PK, KJ, PCM. Involved in management of clinical cohort: JC, EB, PK, KJ, PCM. Wrote the manuscript: EA, PCM, with editorial input from AM, MA. All authors read and approved the final manuscript.

## Acknowledgements

A subset of these data was presented as a poster at the UK Federation of Infection Societies (FIS) conference, November 2014 [28].

## Authors’ information

EB is the lead for the UK STOP-HCV program. PCM is a Wellcome Trust Clinical Research Fellow with interests in chronic HBV and HCV infection.

